# Diverse strains of aster yellows phytoplasma are associated with the potato leafhopper (*Empoasca fabae*) in Eastern Canada

**DOI:** 10.1101/2025.08.28.672916

**Authors:** Ariane Barrot, Joshua Molligan, Abraão Almeida Santos, Nicolas Plante, Edel Pérez Lopez

## Abstract

Phytoplasmas are cell wall–less bacteria that are transmitted by phloem-feeding insects. In Canada, insect vectors of this pathogen are leafhoppers (Hemiptera: Cicadellidae), and they can contribute to significant economic losses. As climate change alters the composition and movement of insect communities, migratory species such as the potato leafhopper (*Empoasca fabae*, Harris 1841), one of the most abundant leafhoppers in Québec, may play an emerging role in phytoplasma diseases. Although *E. fabae* is not currently confirmed to act as a vector, its frequent presence and abundance in fields, along with its potential to acquire phytoplasmas, deserve further investigation. In this study, we tested DNA from *E. fabae* collected in strawberry fields for the presence of ‘Candidatus Phytoplasma’ using PCR, as well as inbred colonies for their ability to transmit this pathogen. The amplicons amplified from positive samples were cloned and sequenced to identify phytoplasma groups and subgroups. Our findings confirmed the presence of multiple Aster Yellows (16SrI-related) phytoplasma strains in *E. fabae*, based on phylogenetic analysis, restriction fragment length polymorphism (RFLP) profiling, and single-nucleotide polymorphism (SNP) profiles. However, the transmission assays did not show vector competence. We propose that although this leafhopper species hosts multiple, possibly new, phytoplasma subgroups, its capacity to transmit the disease remains limited and likely depends on high population density. Overall, these findings emphasize the importance of monitoring common pests like *E. fabae* as indicators of phytoplasma diversity in Eastern Canadian agricultural systems.

## MAIN

Phytoplasmas are wall-less, phloem-limited prokaryotes within the class Mollicutes, provisionally classified under the genus ‘*Candidatus* Phytoplasma’ (Bertaccini et al. 2022). In Canada, a wide range of plant species can host phytoplasmas, along with confirmed leafhopper (Hemiptera: Cicadellidae) vectors (Olivier et al. 2014; Santos et al. 2024). Crops of economic importance at risk of phytoplasma diseases include canola, broad bean, potato, and Canada’s berry industry, which produced approximately 165,000 metric tons in 2024 (Statistics Canada, 2025). As climate warming is expected to increase the presence of leafhopper pests in Eastern Canada (Santos et al., 2025a), understanding which species can act as vectors and the associated phytoplasma strains is a crucial first step in managing phytoplasma diseases.

Since phytoplasmas cannot be cultured *in vitro*, species delineation depends on molecular criteria, notably 16S rRNA gene similarity and restriction-fragment–length polymorphism (RFLP) profiles, which define 16Sr groups and subgroups (e.g., ‘*Ca*. P. asteris’ comprises 16SrI-A, 16SrI-B and so on, now including subgroups classified using double letters like 16SrI-AB). Updated thresholds (>97.5% of 16S operon similarity) together with housekeeping gene comparisons like *tufB, secY, secA, rplV-rpsC*, and *groEL*, can complement in determining species boundaries, in addition to virtual RFLP tools such as *iPhyClassifier* or CpnClassiPhyR (Bertaccini et al. 2022; Muirhead et al. 2019; Wang et al. 2024; Zhao et al. 2009). These tools have helped to identify an increasing number of ‘Ca. Phytoplasma’ species, but vector complexities make surveillance and disease control difficult. For instance, in Canada, 37 leafhopper species have been found to host Aster Yellows phytoplasma (16SrI-related strains), yet only 10 species are competent vectors (Olivier et al. 2014). While not all species are vectors or have demonstrated vector competence, reservoir hosts could enable phytoplasmas to persist in the agroecosystems.

The migratory and polyphagous potato leafhopper, *Empoasca fabae* (Harris) (**Figure 1A**), has been shown to host at least one strain of Aster yellow phytoplasma (Olivier et al. 2014). However, it is still unknown whether phytoplasma-infected individuals are migratory or local and if phytoplasma-infected individuals persist throughout the growing season in Eastern Canada. Additionally, future warming climate trends are expected to increase this species’ range northward in North America (Santos et al. 2025a). Yet, it remains uncertain whether *E. fabae* will expand the range for specific ‘Ca. Phytoplasma’ spp., as available surveys have not yet confirmed *E. fabae* as a vector.

**Figure 1.**
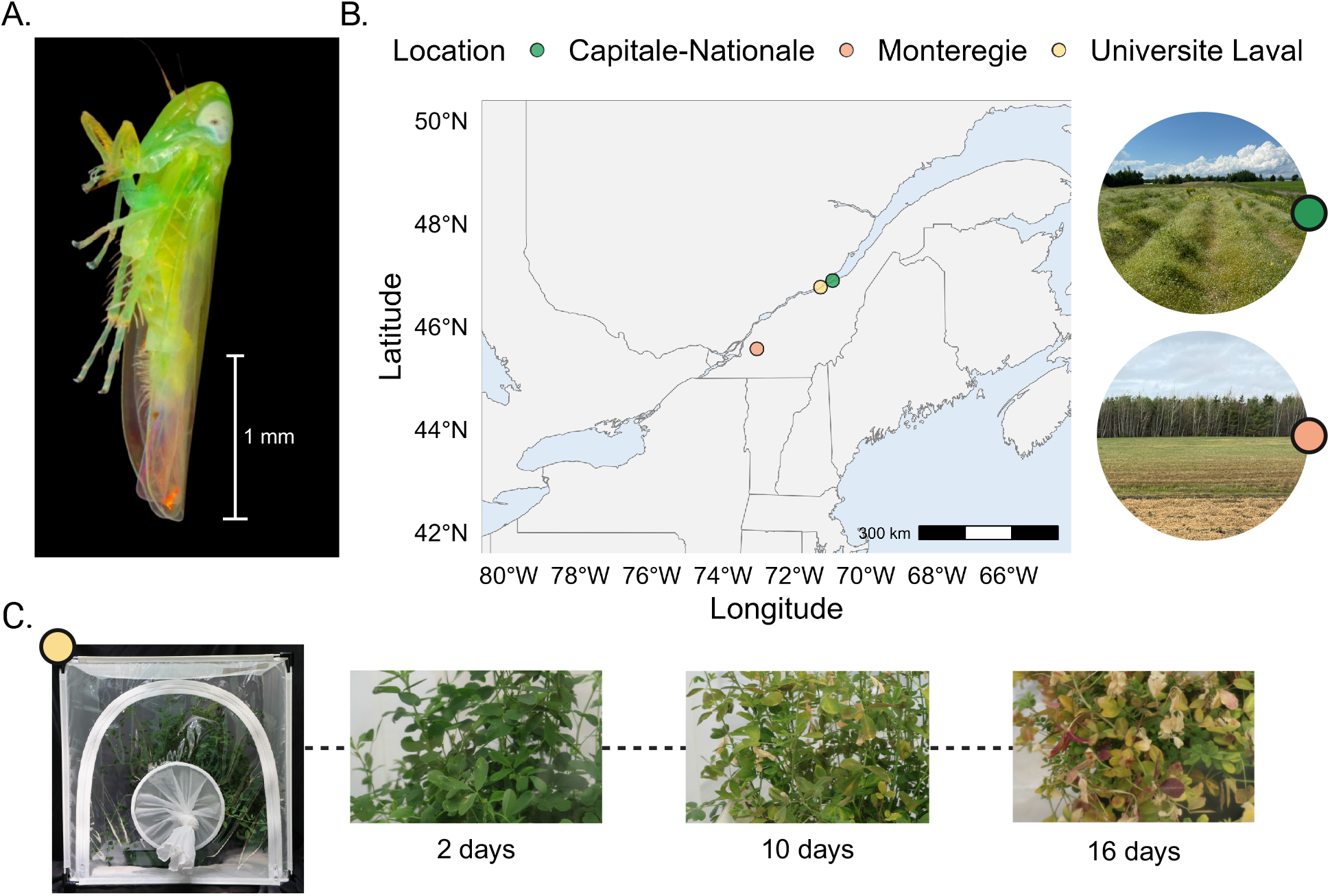
Overview of the model insect, sampling sites, and colony establishment. (**A**) Fixed specimen of *Empoasca fabae* (Photo by Joshua Molligan). (**B**) Map denoting the agricultural collection sites of *E. fabae* and respective images of the sampling locations. (**C**) Greenhouse colonies and depiction of symptoms of hopper burn damage on *Medicago sativa* by *E. fabae* after 2, 10, and 16 days.

Recent citizen-science submissions to Québec’s Plant Protection and Diagnostics Laboratory (LEDP) underscore this province’s growing phytoplasma diversity. Among 34 host plant species reported, five were recorded as new global hosts, and Eastern Canada now records strains related to ‘*Ca*. P. pruni’, ‘*Ca*. P. pyri’, and ‘*Ca*. P. phoenicium’ in blueberries (Brochu et al. 2023). Reports of strawberry green-petal disease, caused by ‘*Ca*. P. asteris’ and ‘*Ca*. P. tritici’ (16SrI group), appear to be rising (Brochu et al. 2023; Plante et al. 2021; 2024; Santos et al. 2025b). This trend coincides with increasing *E. fabae* abundance, as recent field trappings of over 33,000 individuals in Québec strawberry fields identified *E. fabae* as the second most dominant species (Plante et al. 2024).

To further explore the potential of *E. fabae* to host and transmit phytoplasmas, we tested field-caught specimens over two years (2023 to 2024) and established a laboratory colony derived from Eastern Canadian field collections. Here, we report (*i*) screened wild and colony populations of *E. fabae* for phytoplasma presence over the growing season, (*ii*) the characterization of *E. fabae*-associated phytoplasma strains through RFLP and SNP profile, and (*iii*) an assessment of phytoplasma persistence within an inbred colony over time. By characterizing the diversity of aster yellows–related phytoplasmas obtained from *E. fabae* specimens, we suggest that although this species hosts multiple and potentially novel phytoplasma subgroups, its ability to transmit the pathogen remains low under controlled conditions.

The insect specimens used in this study were initially collected from two agricultural regions in southern Québec, Canada, Montérégie (45°35’11.5”N; 73°03’41.5”W) and Capitale-Nationale (46°46’32.4”N; 71°16’48.2”W), as well as from a laboratory colony maintained at Université Laval (**Figure 1B**). All insects tested from the field, and the colony were adults. Field collections in Montérégie were carried out through 2023 using yellow sticky traps, and specimens were preserved in 70% ethanol at 4 °C until processing. In 2024, insects were collected alive from the field in both regions during July using a collection net. The specimens that were not used to extract DNA were used to establish colonies, which were kept under laboratory conditions for further investigation. The cages containing the colonies were kept at 21°C during the day and 18°C at night on a daily cycle of 18:6 hours, while the plant used to feed them was changed every three to four weeks, with weekly fertilization. Laboratory colony specimens were tested between August and December 2024. Additionally, plant tissue samples from alfalfa (Medicago sativa L.), used as the colony food source, were analyzed to assess potential transmission. Samples metadata are summarized in **Table 1**.

**Table 1.**
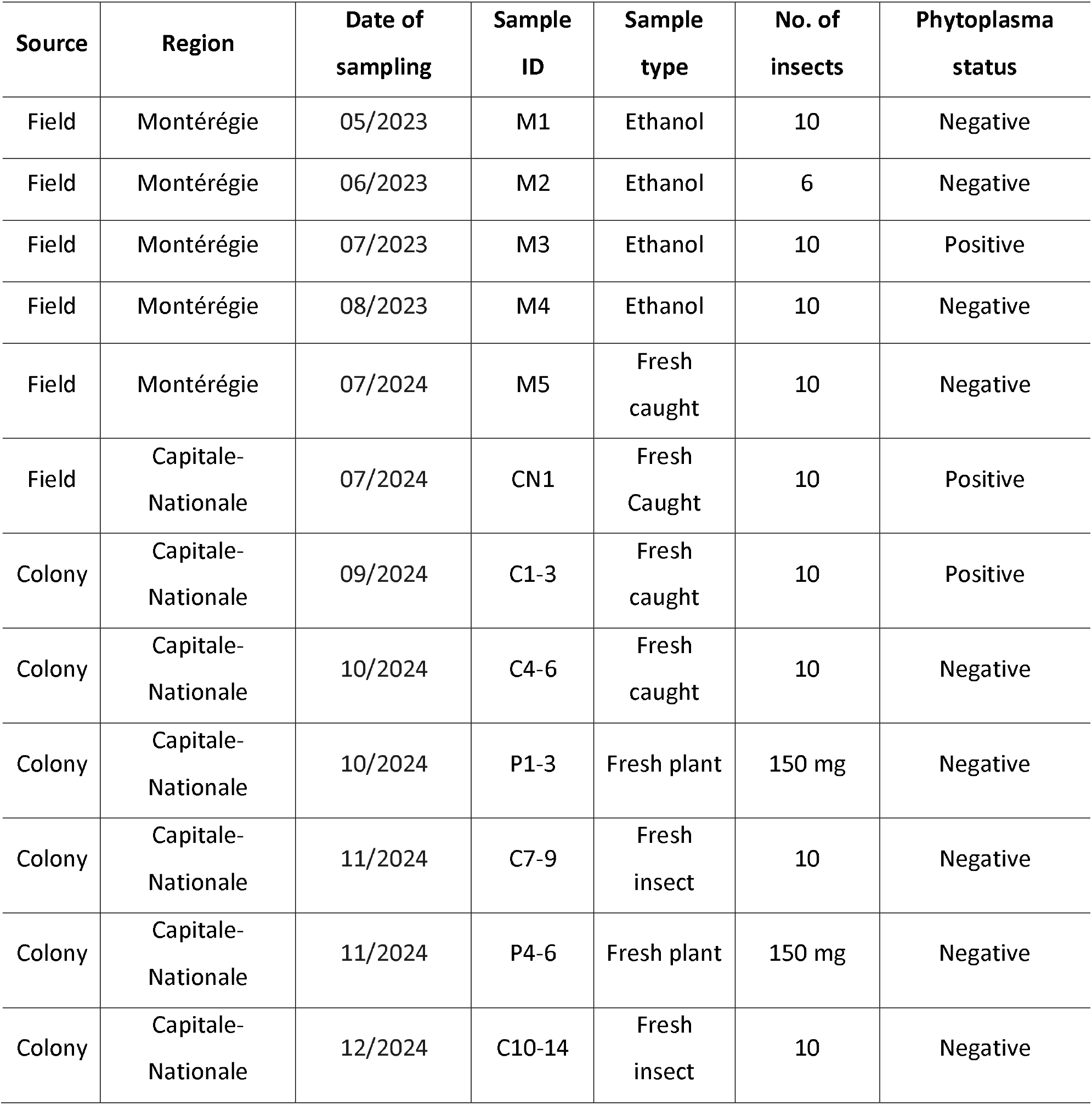
Samples collected from the field and greenhouse, with their location, type, and identification.

Total genomic DNA was extracted from individual insect or plant samples using a CTAB-based protocol as previously (Molligan et al. 2024). Briefly, samples were homogenized in CTAB/PVP lysis buffer, incubated at 65 °C, and subjected to chloroform: isoamyl alcohol (24:1) purification. DNA was precipitated with ice-cold isopropanol, washed with 70% ethanol, and eluted in TE buffer (10 mM Tris-HCl, 0.1 mM EDTA, pH 8.0). DNA quantity and purity were assessed using both a NanoDrop spectrophotometer and a Qubit fluorometer (Thermo Fisher Scientific). DNA integrity was visualized on a 1% agarose gel prepared in TEA 1x buffer and stained with a SYBR Safe DNA gel stain (Thermo Fisher Scientific). Insect and plant samples differed in the initial homogenization method employed. Insects were ground fresh with a mini pestle, whereas plant tissues were first flash frozen with liquid nitrogen before homogenizing with a mortar and pestle.

To determine phytoplasma species identity, a region of the 16S gene was amplified from *E. fabae* gDNA using a nested PCR assay with P1/P7 primers (Deng and Hiruki, 1991; Schneider et al. 1995) followed by a second amplification using R16F2n/R16R2 primers (Gundersen et al. 1994; Lee et al. 1993). Reactions were performed in 50 µl total volume using High-Performance GoTaq® G2 DNA Polymerase in a Ready-to-Use Master Mix (Promega). The PCR assays were carried out under the following conditions for the initial P1/P7 reaction: an initial denaturation step for 30 sec at 95°C, an annealing phase beginning at 65°C for 30 sec, decreasing 1°C for 10 subsequent cycles, and an extension phase at 72°C for 1min 30 sec. Following the initial 10 cycles, the annealing temperature was maintained at 55°C for 35 more cycles. A final extension period at 77°C was maintained for 5min. For the R16F2n/R16R2 reaction, the annealing phase was performed at 68/58°C with all other steps held constant. Binding specificity was influenced by using this touchdown method.

For further subgroup delineation, PCR product was then purified from three representative sampling time points using a PCR purification kit (Qiagen). The purified products were then ligated into the pGEM-T Easy vector (Promega) at a 1:3 vector: insert molar ratio using T4 DNA ligase. The ligation reactions were then transformed into chemically competent *E. coli* TOP10 (lacZΔM15) and plated on LB agar containing ampicillin (100 µg/mL), IPTG (0.5–1 mM), and X-Gal (40 µg/mL) for blue-white screening. After overnight incubation at 37 °C, 25 white colonies were picked into 5 mL LB supplemented with ampicillin and grown overnight. Plasmid DNA was isolated using the QIAprep Spin Miniprep Kit (Qiagen). Insert presence and size were verified by colony PCR, and 20 validated clones from each sample were then sent for Sanger sequencing at the CHUL sequencing facility using universal T7/SP6 primers (Centre hospitalier de l’Université Laval, Québec, Canada).

Cloned amplicon sequences were then trimmed for low-quality bases using respective chromatograms, and aligned using Geneious (Dotmatics, USA, v.2024.0) (Kearse et al. 2012). Species assignment and 16Sr group/subgroup designation were then performed in silico using iPhyClassifier (Zhao et al. 2009). The RFLP patterns obtained were compared with the patterns obtained from reference sequences representative of their respective 16SrI subgroup (**Supplementary Table 1**).

To infer the phylogenic relation of the strains and visualize single-nucleotide polymorphisms (SNPs), the curated set of 16SrI subgroup gene sequences was compared to the reference sequence for each subgroup. First, sequences were aligned in nucleotide mode with MAFFT v7.525 (Katoh and Standley, 2013). The trimmed alignment contained 63 sequences distributed across 1,245 sites, comprising 194 distinct site patterns. This resulting alignment was then subjected to a maximum-likelihood reconstruction with IQ-TREE 2 v2.3.5 (Minh et al. 2020). ModelFinder identified TIM3 + F + G4 as the best-fit substitution model, and branch support was assessed with 1,000 ultrafast bootstrap replicates (**Figure 2A**). To visualize further SNPs, the MAFFT alignment was processed with ciAlign v1.4.1 (Tumescheit et al. 2022), and overhanging, divergent termini were cropped. A similarity-to-consensus plot was then generated to highlight every site that deviated from the modal nucleotide across the final 1,210 bp alignment (**Figure 2A**).

**Figure 2.**
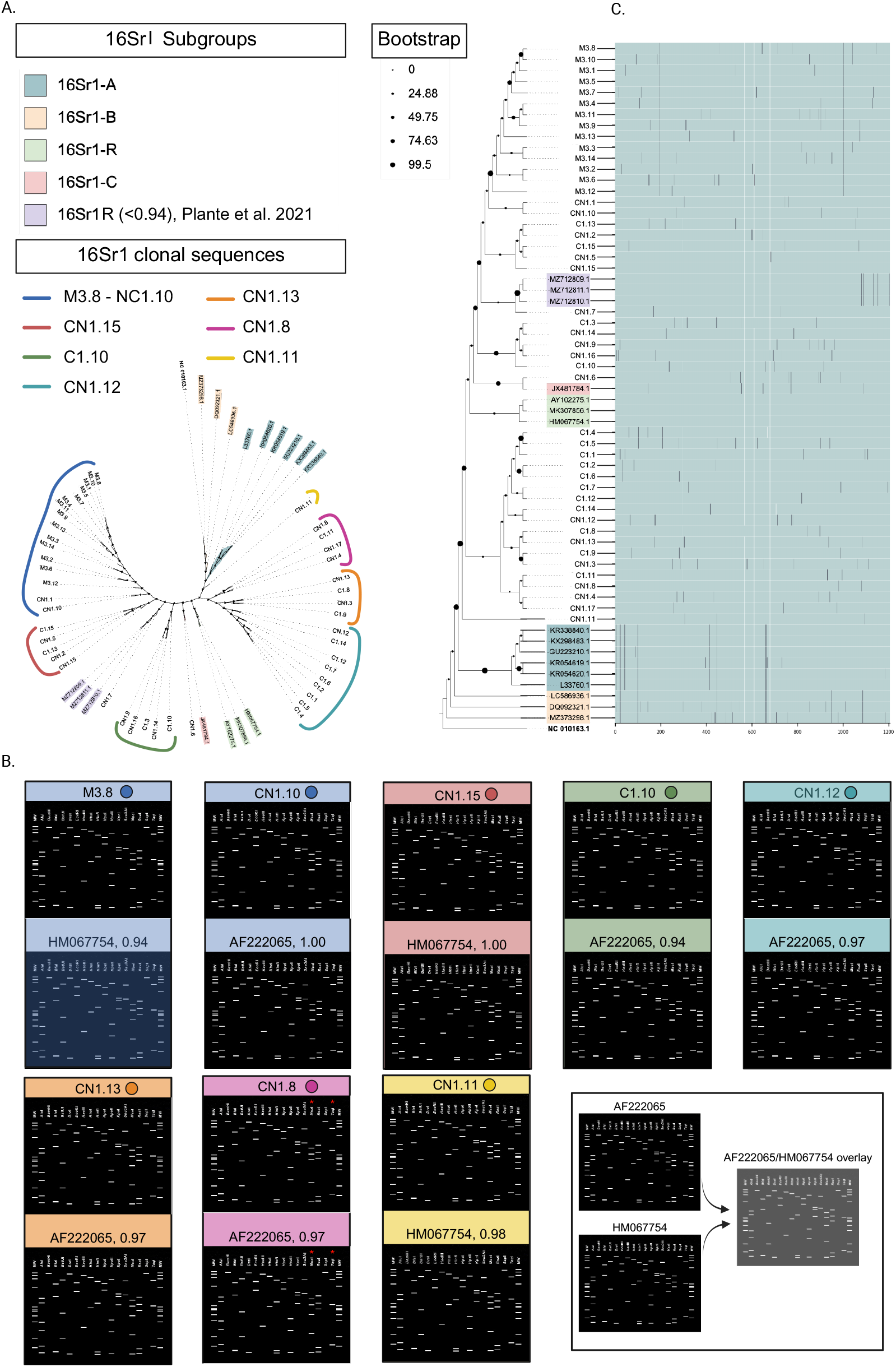
Molecular characterization of phytoplasma strains identified in this study. (**A**) Phylogenetic tree based on the 16S rRNA gene of the samples collected in the field and from the greenhouse, coupled with other selected 16SrI references. (**B**) Virtual RFLP profiles of 16SrI groups from this study based on key restriction enzymes. (**C**) Association with SNPs to highlight the similarities within clusters and phylogenetic tree clades.

Among samples collected in the Montérégie region between May and August 2023, only those from July (M3) tested positive for phytoplasma, whereas samples from May (M1), June (M2), and August (M4) were negative (**Table 1**). After detecting positive samples, we expanded our survey in July 2024 to include both the Montérégie (M5) and Capitale-Nationale (CN1) regions. In this season, phytoplasma was detected only in samples from the Capitale-Nationale site (CN1) (**Table 1**). To evaluate persistence, we established *E. fabae* colonies from both 2024 collections sites. Screening of three colonies in September (C1, C2, C3) confirmed the presence of phytoplasma (**Table 1**). Based on these findings, we next tested whether the pathogen could be transmitted to plant material.

To test phytoplasma transmissibility, three groups of approximately 200 *E. fabae* individuals were randomly assembled from the colony cages (C4, C5, C6) and incubated with three-week-old alfalfa plants (P1, P2, P3) under greenhouse conditions. Fresh plant material and newly transferred insects were sampled initially, followed two weeks later by testing of additional plants (P4, P5, P6) and of colony insects (C7, C8, C9). All insect and plant samples tested negative for phytoplasma at both time points (**Table 1**). Subsequently, insects from five additional cages were screened to further assess pathogen persistence in the inbred colony (C10–C14). These assays confirmed the absence of phytoplasma in later generations (**Table 1**), indicating a lack of persistence within the colony.

To assess the diversity of phytoplasma from field and initial colony samples that were positive, we analyzed 46 16S clones. The RFLP pattern and putative species delineations were obtained with *i*PhyClassifier. Although all the sequences were identified as members of the 16SrI group, we noted two distinct patterns within the ribosomal group that varied in the pattern obtained using the restriction endonucleases *MseI*, and *TaqI* (**Figure 2B**). One specific pattern was present among all clones generated from M3, CN1, and C1, while four clones from CN1 and C1 showed a distinctive RFLP pattern with a 0.97 identity to 16SrI-C reference Clover phyllody phytoplasma (AF222065) (CN1.8, C1.11, CN1.17, CN1.4). Despite the species being the same, ‘*Ca*. P. asteris’, the RFLP pattern differed from all reference sequences, including our previously generated sequences from infected strawberry (*Fragaria* X *ananasa duch*.) (Plante et al. 2021).

Further phylogenetic analysis was conducted with reference members from the 16SrI-A, -B, -C, and -R subgroups including the 46 clone sequences, in addition to SNP visualization. Notably, several of our sequences clustered with established references, including CN1.7 and CN1.6, which grouped with 16SrI-R and 16SrI-C, respectively (**Figure 2C**). The placement of CN1.7 within the 16SrI-R subgroup is surprising, as our earlier work identified this lineage only from infected strawberry plant material. Unlike other 16SrI-R sequences, however, CN1.7 lacked the characteristic SNP sites that consistently define this cluster. In contrast, CN1.6, which grouped with 16SrI-C, carried two distinct SNPs typical of this subgroup. Beyond individual clones, sequences from sample M3 shared conserved SNPs at two positions, reinforcing the observed clustering. Although definitive subgroup assignment of all 46 clonal sequences is not possible because the similarity coefficient for many clones is lower than 0.97, the threshold to erect new subgroups. Together, our results highlight the ability of *E. fabae* to harbor a genetically diverse subset of 16SrI-related subgroups. The phylogenetic analysis presents clusters for all the clonal sequences and 16SrI-subgroup representative sequences. Among them, only CN1.6 and CN1.7 could be grouped within the previously defined subgroups of 16SrI-C and 16SrI-R. At the same time, other sequences appeared to indicate distinctive subclades (**Figure 2C**). When comparing these subclades based on SNPs presence/absence, we observed possible subgroup distinction that RFLP analysis did not allow. Similar methods have been used to designate new subgroups within 16SrXII and 16SrIII and establish phylogenetic relations by incorporating both RFLP and SNP observations (Cheng et al. 2015; Quaglino et al. 2017).

The presence of the phytoplasmas in potato leafhopper over two sampling years indicates that it is present in the field population in July and September and likely to be locally acquired. We also observed that, once the insects were placed in controlled conditions, the presence of phytoplasma was not detected in any of the samples taken from the colony or during the transmission tests. Two scenarios are given to explain these results. First, due to its cell rupture feeding behavior and direct probing, *E. fabae* does not directly inject its saliva into the phloem sieve elements, where phytoplasmas are located (Backus et al. 2005). Thus, phytoplasma acquisition would occur in the presence of an abundant population, allowing the plant to receive a sufficient concentration of phytoplasma to be infected. Such a high-density scenario is noted in Québec from the end of June to the middle of July, where the population peaks and is thought to be composed of local and migratory adults (Jacques et al. 2025; Plante et al. 2024; Santos et al. 2025c). While samples consisting only of migratory individuals were negative (May to June), the positive results occurred when the local adult population was present (from July to September). Nevertheless, this accidental feeding would result in a reduced phytoplasma titer that could lower the likelihood of transmission to healthy plants and offspring, which leads us to the second scenario related to the host plant survival during leafhopper infestation. The alfalfa plants used to feed the *E. fabae* colonies showed hopperburn symptoms, eventually leading to premature senescence within 3 weeks (**Figure 1C**). This may indicate that the plants themselves did not build enough titer to maintain an adequate reservoir of the pathogen for the offspring of *E. fabae* to acquire it. This is reinforced by the role of bacterial concentration inside the insect midgut for transmission to happen. With a relatively quick senescence of the feeding alfalfa host, this could also explain why the pathogen was lost within the colony, as it may not have built enough reservoir within the midguts of *E. fabae*.

In this study, we found that *E. fabae* can host and likely serve as a reservoir for multiple phytoplasma Aster Yellow subgroups in local populations in Québec, although we did not detect transmission to host plants. This suggests that crop pests can serve as valuable indicators of pathogen diversity in the field, even though they may not serve as vectors. Regular screening of abundant pests such as *E. fabae* could help focus surveillance and guide management practices for potentially devastating ‘*Ca*. Phytoplasma’ spp. We suggest that further work should test if, and under what conditions, *E. fabae* might serve as a vector, and to validate further the distinctive lineages that this species carries in Eastern Canada.

## DATA AVAILABILITY STATEMENT

All sequences generated in this study are deposited under the GenBank Accession Numbers PX238692 to PX238737.

## AUTHOR CONTRIBUTIONS

Conceptualization, J.M. and E.P.L.; Investigation and lab work, A.B. N.P., and J.M.; Cloning, J.M.; Visualization, J.M. and A.A.S.; Resources (specimen photography), J.M.; Writing – original draft, A.B. and J.M.; Writing – review & editing, A.B., J.M., N.P., A.A.S., and E.P.L.; Supervision & Resources, E.P.L.; A.B. and J.M. contributed equally. E.P.L. is the corresponding author.

## ACKNOWLEDGEMENTS

We thank Anne-Sophie Brochu for her laboratory expertise and assistance, as well as Jordanne Jacques for sample collection and specimen identification.

