## Supplementary material for "Diverse strains of aster yellows phytoplasma are associated with the potato leafhopper (*Empoasca fabae*) in Eastern Canada": Table S1

**SUPPLEMENTARY TABLES**

**Supplementary Table S1.** List of the sequences used from the literature with their accession number, species, and the reference to the source article

| **Grouping** | **Accession number** | **Species** | **Reference** |
| --- | --- | --- | --- |
| 16SrI-A | GU223210.1 | Phytoplasma asteris | (Jomantiene et al., 2010) |
| 16SrI-A | KX298483.1 | Phytoplasma asteris | (Ivanauskas et al., 2016) |
| 16SrI-A | KR338840.1 | Phytoplasma asteris | (Ivanauskas et al., 2016) |
| 16SrI-A | KR054620.1 | Phytoplasma asteris | (Valiunas et al., 2015) |
| 16SrI-A | KR054619.1 | Phytoplasma asteris | (Valiunas et al., 2015) |
| 16SrI-A | L33760 | Phytoplasma tritici | (Lee et al., 1993) |
| 16SrI-B | DQ092321.1 | Phytoplasma asteris | (Santos-Cervantes et al., 2008) |
| 16SrI-B | MZ373298.1 | Phytoplasma asteris | (Tseng et al., 2022) |
| 16SrI-B | LC586936.1 | Phytoplasma asteris | (Jonson et al., 2020) |
| 16SrI-C | JX481784.1 | Phytoplasma tritici | (Zhang et al., 2013) |
| 16SrI-R | AY102275 | Phytoplasma tritici | (Jomantiene et al., 2002) |
| 16SrI-R | MK307856.1 | Phytoplasma asteris | (Babaei et al., 2021) |
| 16SrI-R | HM067754.1 | Phytoplasma asteris | (Jomantiene et al., 2011) |
| 16SrI | MZ712809.1 | Phytoplasma asteris | (Plante et al., 2021) |
| 16SrI | MZ712810.1 | Phytoplasma asteris | (Plante et al., 2021) |
| 16SrI | MZ712811.1 | Phytoplasma asteris | (Plante et al., 2021) |
